# Thymic Rejuvenation by Adult Grafts in the Lymph Node

**DOI:** 10.1101/2022.06.06.495022

**Authors:** Aparna Rao, Avni Gupta, Eric Lagasse

## Abstract

Immunologic aging is defined as a process in which the immune system undergoes tremendous decline in T cell numbers and function, thus affecting one’s ability to effectively mount an immune response. The degeneration of the thymus, the primary immune organ responsible for T cell development, is central to immunologic aging. Various studies have assessed thymic regeneration as an effective way to offset immune function decline. In this study, we determined the effect of mouse adult thymus transplants in the lymph node of athymic nude mice to rejuvenate thymus function. We transplanted thymuses of increasing ages and assessed engrafted ectopic thymuses by histology as well as for blood T cell numbers and function. We observed that transplanting aged thymuses, up to 8 months old, in the lymph node regenerated thymus function and corresponding T cell activation with some thymic rejuvenation when compared to the expected native thymus in control animals. However, transplanting 11- and 14-month old thymus in the lymph node had decreased-to-limited thymic function with no thymic rejuvenation detected. These observations provide important insights into the plasticity and regenerative capabilities of the thymus during age-associated involution.

## Introduction

The thymus is a central organ of the immune system, necessary for T cell development and maturation [1]. The thymus is crucial for education of T cells, and a dysfunction or loss of thymic function can result in a variety of disorders, including autoimmunity, graft versus host disease and loss of immune activity, as seen in DiGeorge syndrome [2]. Therefore, restoration of thymus function can provide considerable benefit to individuals whose T cell immune functions have been decimated in various settings. As we age, the thymus is also one of the earliest organs to undergo a decline in function, a process called ‘thymic involution’ [3, 4]. Thymic involution is characterized by a decrease in thymic epithelial cell coverage, accompanied by a decrease in T cell output. This reduction in T cell development concurrent with a decline in naïve T-cells and less diverse TCR repertoire lead to a weak adaptive immune response during infections [5]. Sometimes, a defective thymic education process also precedes to the release of a higher number of autoreactive T cells to the periphery, thus increasing the susceptibility to autoimmune disorders [6, 7].

Our group has extensively studied the use of the lymph node as a site of ectopic organogenesis. Lymph nodes are highly vascularized sites with central access to blood vessels, the lymphatic system and serve as spatially highly organized hubs of immune cell interactions [8]. In addition, lymph nodes are capable of facilitating the migration and proliferation of immune cells, but also in some cases, the migration and proliferation of metastasizing epithelial tumor cells [9]. Earlier, we have demonstrated that transplanting a neonatal thymus in the lymph node was able to reconstitute the immune system in athymic nude mice [10]. The lymph node also provides a more clinically relevant site for transplantation compared to other sites such as the subcapsular space of the kidney capsule previously published [7, 11]. While thymus transplant in the lymph node could technically be a feasible approach for thymus regeneration, our earlier approach using neonatal thymuses had limited potential and therefore, we decided to replicate our earlier efforts to reconstitute the immune system in nude mice by transplanting older thymus into lymph node.

We wanted to primarily evaluate whether transplanting an adult and aged thymus in an enriched environment such as the lymph node will regenerate thymic function comparable to that of an age match native thymus control. For this purpose, we transplanted thymus of different ages in the lymph node of 6 weeks old nude mice. The ectopic thymuses were then analyzed for engraftment, molecular signatures, and T cell function. We observed that thymuses that were up to 8 months old were able to sufficiently engraft in the lymph node and reconstitute immune function in nude mice with a molecular signature representing a significant delay in the expression of important aging factors. However, thymuses 11 and 14 months old were not able to perform in a similar manner. The limited function of these older thymuses engrafted in the lymph node translated into decreased T cell development, decreased circulating blood T cells and eventually the deficiency of a T-cell immune response in mice. We believe these results provide important insights into the decrease in plasticity of thymus along with progressing age, and may lead to novel interventional strategies to reverse thymic involution.

## Materials and Methods

### Mice and tissues

6 weeks old athymic nude mice (NU/J, Jackson Laboratory) were used for thymus transplants. Donor thymic tissues were isolated from Balb/c-GFP mice of different ages. Age was calculated according to date of birth. Donors and recipients were not matched according to gender. Blood collection (100μL) was performed using the submandibular bleeding technique. Mice were bred and housed in the Division of Laboratory Animal Resources facility at the University of Pittsburgh Center for Biotechnology and Bioengineering. Experimental protocols followed US National Institutes of Health guidelines for animal care and were approved by the Institutional Animal Care and Use Committee at the University of Pittsburgh.

### Thymic transplantation

Thymuses were harvested from BALB/c-GFP+ transgenic mice of different ages. Age was calculated according to date of birth. Thymus was isolated from the thoracic cavity and minced to get 1-2mm pieces. The pieces were vortexed vigorously and treated with 2-deoxyguanosine in order to get rid of T cells, as described elsewhere [12, 13]. The jejunal lymph nodes of recipient mice were exposed, and minced thymic tissue was injected through a 20G needle. Light cauterization was used to seal the opening. The laparotomy was closed with surgical sutures. One donor thymus was injected per recipient mouse.

### Tail skin graft

Recipient mice (athymic nude mice) containing an ectopic thymus (BALB /c-GFP+) in the lymph node were anesthetized, and the tail skin (5 mm × 5 mm) from BALB /c or C57BL/6 mice was grafted on the left and right lateral sides of the superior dorsal region of the recipient mouse, respectively. A bandage was applied and removed 7 days after surgery. The grafts were observed every other day for rejection. A graft was considered as accepted if the skin was on the graft more than two weeks after the transplant. Grafted skin that necrosed and wilted was considered as rejected.

### Antibodies

Antibodies specific to the following antigens were purchased for immunohistochemistry: keratin 5 (Covance), keratin 8 (DSHB), and Ki67 (Abcam). Antibodies specific to the following antigens were purchased for flow cytometric analysis: APC mouse CD3-ε, APC-Cy7 mouse CD8-α, phycoerythrin (PE)-Cy7 mouse CD45, PE mouse CD4, PE mouse FoxP3 and BV421 mouse CD25 (BD Biosciences). Appropriate isotype control antibodies (BD Biosciences) were used to estimate background fluorescence.

### Immunohistochemistry

Tissue was fixed in 4% paraformaldehyde for 4 h, stored in 30% sucrose for 12h and then embedded in optimal cutting temperature (OCT) medium, frozen and stored at −80 °C. Sections 5-7μM were mounted on glass slides and fixed in cold acetone for 10 min. For immunohistochemical staining, sections were washed with PBS and blocked with 5% bovine serum albumin (BSA) or milk for 30 min. Sections were then incubated with primary antibody for 1 h and secondary antibody for another 1h. Sections were mounted with mounting medium containing Hoechst. Images were captured with an Olympus IX71 inverted microscope.

### Image analysis

Entire frozen sections stained with relevant antibodies (CK5/CK8/ Ki67) were imaged using a scanning microscope (Nikon90i) and all the images were stitched together to form one composite image using the Nikon Elements software. Analysis was performed using the Nikon elements software. The image was separated into separate channels. A thresholding mask was created to identify all positive signals and exclude negative signals. Total number of cells was calculated by analyzing the number of DAPI+ cells. % Engraftment was calculated as follows – (Number of GFP^+^ cells/Total cells) x100. %CK5/CK8/Ki67 was calculated as follows: (Number of GFP^+^ surface marker^+^ cells/Total cells) x100.

### Quantitative PCR Analysis

Native and ectopic thymuses were excised and RNA was isolated using the RNeasy kit (QIAGEN). RNA was quantified with the Nanodrop 2000c Spectrophotometer (Thermo Scientific). Reverse transcription was performed using iScript RT Supermix (Bio-Rad) followed by real-time PCR on an ABI StepOne Plus Real-Time PCR machine (Applied Biosystems). TaqMan probes and primers were purchased from Life Technologies (***Table S1***). Expression was normalized to *Gapdh* as well as expression levels of individual genes in a non-transplanted lymph node. Error bars represent upper and lower error limits based on replicate variability observed in triplicates.

### Flow cytometry

Whole blood was collected in K2EDTA collection tubes (Terumo Medical). One hundred microliters of blood were added to cold fluorescence-activated cell sorting (FACS) tubes. Three milliliters of Red Blood Cell Lysing Buffer (Sigma) was added to each tube, lightly vortexed and incubated for 5 min. Two milliliters of flow buffer (2% FBS in HBSS) was added to the tubes, mixed and centrifuged at 500*g* for 5 min. Antibodies were added at a dilution of 1/50 to the cells and mixed by gentle pipetting. Reactions were incubated in the dark in an ice slurry bath for 30 min. For FoxP3 staining, cells were permeabilized with eBioscience Fix/Perm Buffer and then stained for 1hr with anti-FoxP3-PE antibody (eBioscience). After a final wash, permeabilized cell pellets were resuspended in 400μl of flow buffer and live cell pellets were resuspended in 400μl of flow buffer containing the viability dye Sytox Blue (Life Technologies). Cells were analyzed using a Miltenyi MACSQuant Analyzer. Live cells (Sytox Blue negative) were used for subsequent analysis of relevant populations. For Tregs analysis, total cells present in the sample were analyzed.

### T cell stimulation assays

#### Preparation of T cells

Responder T cells (CD3+) were purified from splenocytes of thymus-transplanted or IS control mice by MACS. Celltrace™ CFSE (Molecular Probes, Life Technologies) was dissolved in DMSO as 10 mM stock solutions (stored at − 20 °C). Cells were labeled with CFSE dye as previously described [14]. Briefly, lymphocytes were resuspended to 1 × 107/mL in PBS (Life Technologies) and a final concentration of 10 μM CFSE dye added to 1 mL aliquots of lymphocytes, vortexing immediately to ensure rapid and homogeneous labeling of cells. Cells were incubated at 37°C for 15 min, then washed 3 times with RPMI 1640 supplemented with 10% FBS. *Culture conditions.* CFSE-labeled T cells were re-suspended at a concentration of 1 × 10^6^/mL each in RPMI 1640 supplemented with 10% FBS, 10 mM HEPES, and 1 mM L-glutamine, at a final volume of 200 μL/well in 96 well U-bottomed plates (Nunc). Anti-CD3 stimulating antibody (BD Biosciences) was added to each well at a final concentration of 5μg/ml. Cells were cultured at 37 °C in 5% CO2 for 72 hours and analyzed by flow cytometry. The gating strategy used to analyze the cells was as follows: Live > CD45+/CD3+ > Celltrace™. Cell-free culture supernatants were assessed for IFNγ levels by ELISA (R&D Systems).

### Statistical analyses

Statistical significance was determined with an unpaired two-tailed Student’s *t*-test for the data shown in indicated figures.

## Results

### Isolation and transplantation of thymus

Thymuses were isolated from GFP+ BALB/c mice of different ages (neonatal day 2 to 4, and 6 weeks, 3 months, 6 months, 8 months, 11 months, 14 months old thymuses). Various studies have demonstrated the decrease in epithelial cell coverage in thymus with increasing age [15–17] and using qualitative as well as quantitative immunohistochemical techniques, we were able to replicate these findings (***Figure S1***). To assess whether engraftment and function of an ectopic thymus in the lymph node was affected by age, we transplanted different aged thymuses in young 6-week-old adult nude mice LNs. Thymuses were previously treated with 2-deoxyguanosine to eliminate T cells and then transplanted with a single thymus in the jejunal lymph node. Thymuses were GFP+ in order to differentiate between donor and host tissue. 6 weeks after transplant, the ectopic thymuses were analyzed for engraftment by determining number of GFP+CK5+/8+ cells, respectively (***Figure 1A***). Percentage of engraftment was determined by looking at the proportion of GFP+ cells in the entire section. By analyzing tissue sections among the different groups, we observed an inverse correlation between age of the thymus and the engrafted tissue, an observation expected due to the decreasing size of the tissue transplanted with the increasing age of the thymuses (***Figure 1B***). In particular, the proportion of thymic cortical epithelial cells (cTECs, CK8+ cells) engrafted in the lymph node were negatively affected by the transplantation process, when compared to medullary thymic epithelial cell (mTECs, CK5+ cells). This difference in engraftment was not present for 11- and 14-week-old thymuses, where less tissue appears to engraft (***Figure 1C***). Furthermore, cell proliferation in transplanted thymuses (GFP+ Ki67+) was significantly reduced in these older thymuses, possibly causing an additional decrease of cell coverage (***Figure 1D***).

**Figure 1:**
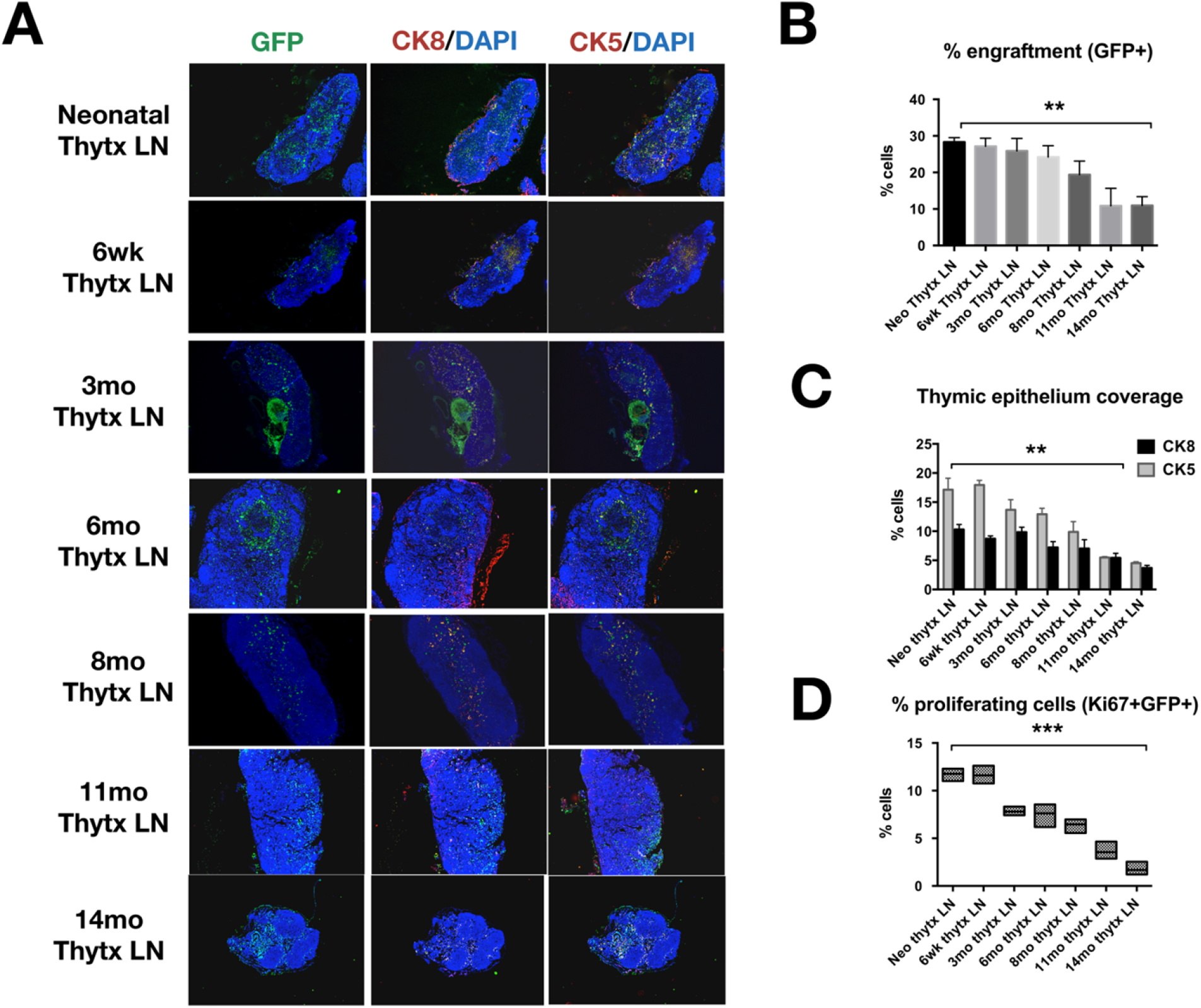
Engraftment of thymuses of different ages in the LN of nude mice. BALB/c GFP thymuses from different aged mice (as indicated) were transplanted (Thytx) into the lymph node (LN) of nude mice. 6 weeks after transplant, the transplanted LNs were isolated and analyzed. A) 5-7 μm frozen sections of LNs transplanted with different aged GFP+ thymuses were assessed for thymus engraftment (GFP+, green, left panel), cortical epithelial cells (Cytokeratin 8, CK8, stained red, middle panel) and medullary epithelial cells (Cytokeratin 5, CK5, stained red, right panel). DAPI was used to counterstain nuclei. Scale bar = 200 μm. B), C) and D) The entire frozen section was imaged using a scanning fluorescence microscope (Nikon 90i) and assessed for various markers. Images were then stitched together and analyzed using the Nikon elements software. B) Percentage of engraftment of transplanted thymus was calculated using the formula: % engraftment = (Number of GFP^+^ cells/Number of DAPI+ cells) x100. C) Cortical and medullary epithelial cell coverage was calculated using the formula: % epithelial cell coverage = (Number of GFP+CK5/CK8+ cells/ Number of DAPI+ cells) x100. D) Percentage of proliferating cells was calculated by % proliferating cells = (Number of GFP+ Ki67+ cells/ Number of DAPI+ cells) x100. **P<0.005, ***P<0.0001

### Improvement in expression of transcripts involved in thymic functions from ectopic thymus

Thymus involution is associated with changes in expression of multiple transcripts important for cells development and functions. Therefore, we compared the expression profile of transcripts differently expressed during aging in both native and ectopic thymuses. We performed quantitative real time PCR for CXCL12 and its receptor CXCR4, CCL25, DLL4, FoxN1 and IL7 (***Figure 2***). Both CXCL12, and CCL25 are important regulators of T cell recruitment and development. Theses transcripts were down-regulated in aging of native thymuses, beginning as early as 3 months [18, 19]. CXCR4 expression on the other hand, began to increase as the age of the thymuses increased. Markers of thymic function such as DLL4 and FoxN1 were also significantly down-regulated in aged mice (***Figure 2***). These results are concordant with observations made by other research groups in the field [20–22]. Interestingly, we observed that transplanting aging thymuses into lymph nodes suggest a delay in aging with an increase in the presence of transcripts expression generally altered in the course of thymic involution (***Figure 2***). Yet, transplantation of 11- or 14-month-old thymuses in lymph node did not have a significant impact on expression of the genes assessed, and were similar to those observed in age-matched native thymuses (***Figure 2***).

**Figure 2:**
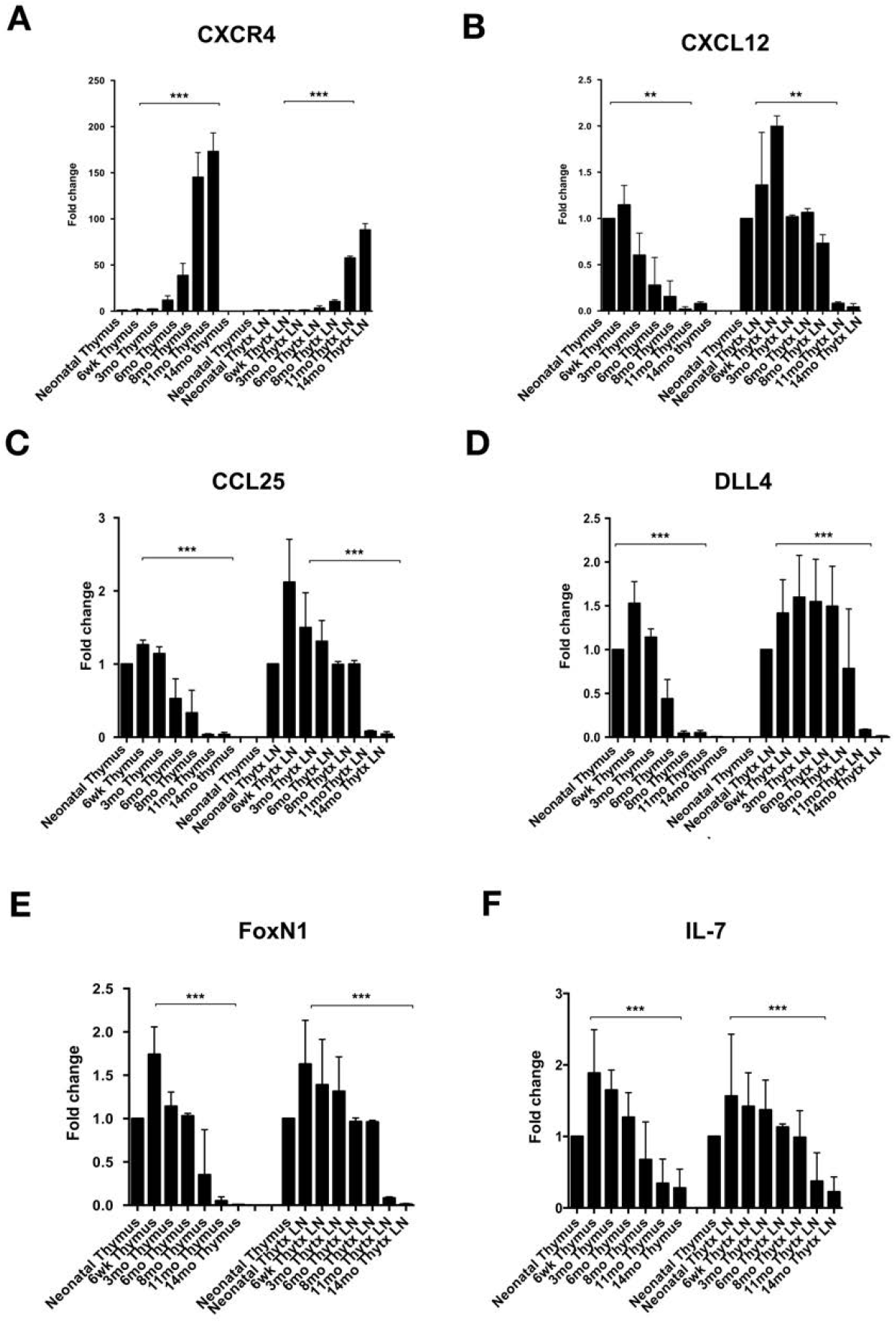
Molecular analysis of aged thymuses, native and transplanted into LNs. Native thymuses of different ages from BALB /c mice as well as aged match thymuses transplanted in Nude mice LNs were isolated and mRNA was extracted from them. The samples were then analyzed for comparative gene expression of several genes (as indicated) by quantitative real-time PCR. Genes assessed for expression were CXCR4 (panel A), CXCL12 (panel B), CCL25 (panel C), DLL4 (panel D), FoxN1 (panel E) and IL-7 (panel F). Neonatal thymus was used as a reference control for all the other samples, and levels were normalized to expression levels form a non-transplanted lymph node. **P< 0.005, ***P<0.001

### Reconstitution of T-cell populations after adult thymus transplantation into lymph node

One of the major repercussions of thymus age-associated involution is the decrease in T cell output, as supported by decreasing proportion of CD4+CD8+ T cells present in the thymus [23–25] (***Figure S2***). In general, studies have delineated the decrease in thymic function [26]. Here, we assessed the correlation between the age of the thymus being transplanted and the extent of T-cells reconstitution in recipient nude mice by analyzing both blood and thymus derived T-cells. For this purpose, we transplanted nude mice lymph node with thymuses of different ages, as described earlier. Six weeks after transplant, mice were bled in order to determine the proportion of circulating T cells in the periphery. Since the athymic nude mice have negligible to no T cell numbers inherently due to the lack of thymus, all the T cells observed in the periphery were assumed to have been generated by the ectopic thymus and as previously showed with transplantation of newborn thymus [10]. Furthermore, transplanted thymuses were derived from GFP+ mice, depleted for T cells by 2-deoxyguanosine treatment and the absence of GFP+ T cells in the blood was confirmed by flow cytometry (not shown). The amount of total CD3+ T cells in the blood was inversely proportional to the age of the transplanted thymus (***Figure 3B and 3C***). The proportion of CD4+ T cells among CD3+ T cells remained fairly constant (***Figure 3B and 3D***). On the other hand, with increasing age of the transplanted thymus, there was a 4-fold reduction in CD3/CD8+ T cell numbers (***Figure 3B and 3E***). As documented previously, while the CD4 T-cell population is relatively stable during aging, the CD8 T-cell population undergoes more drastic changes suggesting that CD4 T cells are more resilient to resist age-associated changes [11]. Interestingly, the proportion of regulatory T cells (Tregs) also seemed to be negatively affected as the age of the transplanted thymus increased (***Figure 3B and 3F***). The majority of Treg production occurs early after birth (within the first week) and declines with age [27, 28]. About 10 weeks after transplant, we harvested the ectopic thymuses and analyzed them for their ability to promote T cell development with the proportion of CD4/CD8 double-positive T cells present. Transplanted thymus into the lymph node was able to restore T-cell development, particularly with thymuses up to 8 months of age. Even older thymuses transplant (11 and 14 months) had restoration of some T-cell differentiation in nude mice, although at much lower levels (***Figure 3A***) particularly when compared to native thymus age match controls (***Figure S2***). In summary, our findings indicate that the lymph node is an efficient site to support engraftment of thymus and the development of T-cells, even during the process of involution.

**Figure 3:**
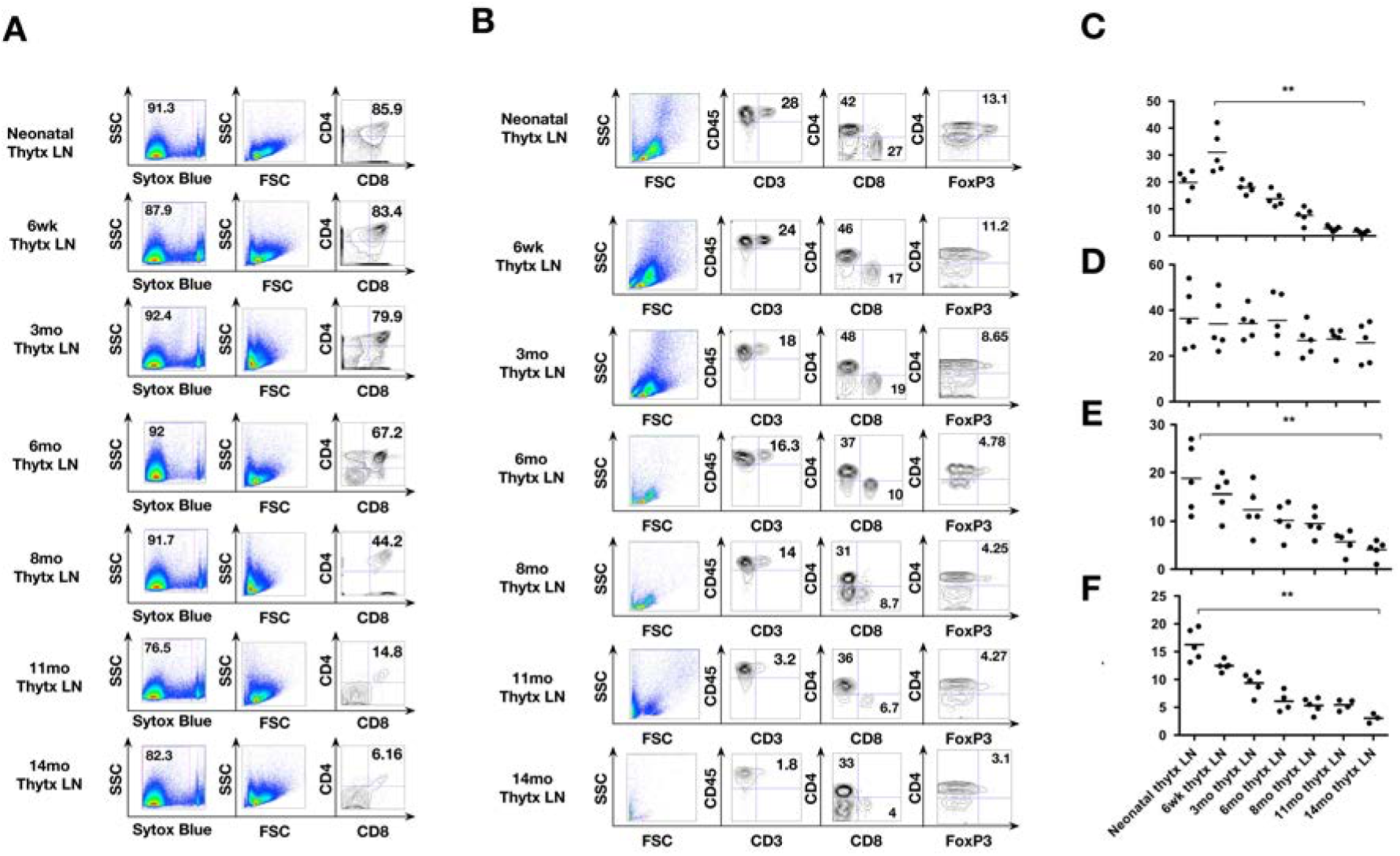
Ability of transplanted thymuses to reconstitute the immune system in nude mice. Nude mice transplanted with thymuses of different ages were bled 6 weeks after transplant. Blood cells were analyzed for various T cell subsets by flow cytometry. A) Thymuses from mice of different ages were harvested and dissociated into single cell suspensions. The cells were analyzed for double-positive CD4/CD8 T cells. Plots shown are representative of tissues from one mouse from a cohort of 3-5 mice in each group. Gating strategy used was as follows: SSC vs. Live (Sytox Blue) →FSC vs. SSC →CD4 vs. CD8. B) Representative flow cytometry plots of bleeds from all groups. Plots shown are representative of bleeds from one mouse from a cohort of n=5 mice in each group. Gating strategy used was as follows: SSC vs. Live (Sytox Blue) → FSC vs. SSC → CD45 vs. CD3 → CD4 vs. CD8 → CD4 vs. FoxP3. Summary graphs of the proportion of CD3+ T cells (B) CD3+CD4+ cells (C) CD3+CD8+ cells (D) and CD4+FoxP3+ cells (E) obtained by flow cytometry from mice in all cohorts. **P< 0.005.

### T-cell function from aging ectopic thymus in a lymph node

As we age, the T cells produced by thymus have a decreased ability to combat infections and neoplasms. We tested the functional capabilities of the T cells produced by the ectopic thymuses in all cohorts using the following read-outs: T cell proliferation using CFSE dilution, an IFNγ assays, and allogeneic skin graft rejection. CD3+ T cells from the spleen of thymus-transplanted mice were labeled with Cell-Trace™ CFSE and stimulated with anti-CD3 antibody. T cell stimulation, as evidenced by CFSE dilution and IFNγ levels (***Figure 4A and 4B***, respectively), were significantly higher in mice transplanted with thymuses up to 8 months old, versus older thymuses. These results correlate well with T cell numbers in the host and productivity of the ectopic thymus in general, and age match T cells from control mice (***Figure S3***). Furthermore, we performed fully allogeneic tail skin grafts (C57BL6/J) on the flanks of these mice and monitored them for allogeneic rejection. Skin is highly immunogenic and therefore elicits a strong immune reaction, even if total T cell numbers are low. However, corresponding to the T cell stimulation assay results, we observed significantly quicker rejection of all allogeneic skin grafts in mice transplanted with thymuses that were 8 months old or younger (100% rejection rate). Mice with older thymus transplants (11 month and 14 months) had just a 20% skin graft rejection rate (1/5 grafts rejected). These results are reflected in the corresponding circulating T cells numbers observed (***Figure 3***), and their proliferation index (***Figure 4A***), which might have played a major role in graft rejection.

**Figure 4:**
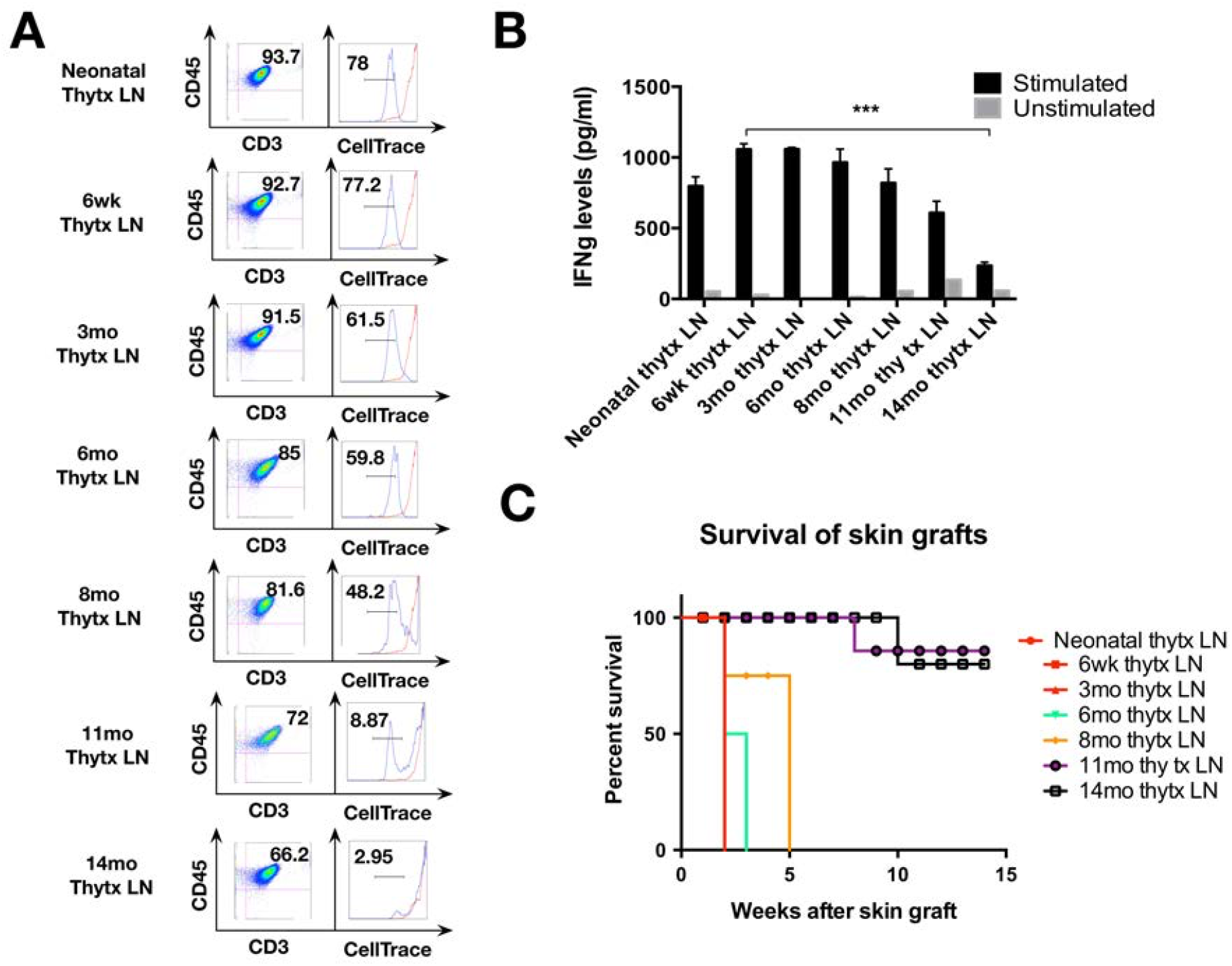
Functional efficacy of T cells generated by transplanted thymuses. A) Splenocytes were isolated from thymus-transplanted mice; CD3+ cells were isolated by MACS and labeled with CellTrace™. The cells were then stimulated with anti-CD3 antibody and the cells were assessed for Celltrace™ dilution (Panel A, blue histograms). Unstimulated cells were used as controls (Panel A, red histograms). B) IFNγ levels from supernatants of cell cultures in (A). ***P<0.0001. C) Thymus-transplanted mice were grafted with C57BL/6 tail skin on their flanks. The grafts were then monitored for rejection every week. Skin grafts were considered as rejected when the grafted skin wilted and fell off the graft pad. (n=3 to 5 mice per group).

## Discussion

We have previously established the lymph node as a viable surrogate site for thymus regeneration and function using newborn thymus [10]. In this study, we evaluated the ability of different aged adult thymuses to reconstitute the immune system of nude mice with the lymph node as a site of transplantation. The ectopic thymuses (thymuses transplanted into LNs) were then analyzed for engraftment, specific molecular signatures and histology. In addition, the transplanted mice were also assessed for blood T cell numbers and function.

Our results recapitulate and extend earlier reports that the generation of functional T-cells by grafted thymus tissues decreases with the age of the graft and suggest that the extent to which T cells can mature is dependent on the degree of thymic involution [7]. However, generally aging also affects various stages of T cell development, from age-related alterations in T-cell progenitor cells and disruption in thymic microenvironment niche, to the changes in key signaling molecules that modulate thymopoiesis and the subsequent decline in T cell output and immune functions [29].

Here, we showed an improvement in expression of certain transcripts involved in these thymic functions from engrafted ectopic thymus. CXCL12 and its receptor CXCR4 are important chemokines involved in numerous biological processes during T-cell development in the thymus. CXCR4 increases dramatically at a later stage of thymus involution while CXCL12 decreases. Engrafted and involuted ectopic thymuses showed a transcriptional decrease in CXCR4 expression while an increase in CXCL12 expression when compared to aged matched native thymuses. Similarly, chemokine ligand 25 (CCL25) and CXCL12 are critical for recruitment and homing of lymphoid progenitors in thymus and aging thymus displayed a marked reduction in CXCL12 and CCL25 mRNA expression during thymus aging process. Engrafted and involuted ectopic thymuses increased expression of CCL25 and CXCL12 similarly to what was reported in an early study on calorie restriction and age-related thymic involution [30]. Delta-like 4 (DLL4) is indispensable in thymic environment specific for T cell development [31]. DLL4 transcript in involuted ectopic thymuses was generally increased up to 8 months. One master transcription factor for thymic epithelial development program is fork-head box protein-N1 (FOXN1)[22, 32], which has been shown to modulate TEC patterning in the fetal stage and TEC homeostasis in the post-natal thymus. One striking example of FOXN1 function is in the athymic nude mice with spontaneous mutation of *Foxn1*. FoxN1 transcriptional level was increased in ectopic thymuses particularly at 8 months of age. Finally, Interleukin 7 (IL7) is produced by stromal cells in thymus, and promotes both survival and differentiation of immature, and mature single positive thymocytes [33]. IL7 is expressed in the thymic microenvironment and its downregulation is associated with thymus involution. An increase in IL7 transcripts was detected particularly in transplanted thymuses 6 and 8 months old. In summary, many of the gene’s transcripts levels affected by age related thymic involution and up to the age of 8 months were reversing their expression profiles after thymus transplantation into the mouse lymph node. This result suggests limited rejuvenation of aged involuted thymus by factors extrinsic to the thymus. A similar observation was previously described by Yamada et al. [34] in an allogeneic vascularized transplant setting with the miniature swine model. In this model, 20 months old involuted thymuses were transplanted into juvenile 4 months old animals. These aged involuted grafts underwent thymic rejuvenation, suggesting that thymic involution can be controlled by the host environment [34]. In our study, we transplanted 6 weeks old mice which are considered as young adult mice and could have provided the environment necessary for thymus rejuvenation. In addition, our mice involuted thymuses transplantation was ranging from neonatal to adult 14 months old mice. Interestingly, both 11- and 14-month-old thymuses presenting no rejuvenating activity after transplantation by transcriptional analyses. This observation with older thymuses transplant has not been reported by Yamada et al., possibly because 20-month old involuted pig thymuses transplanted in their study are still considered as young adult with epiphyseal lines just closing at this age for miniature pigs [35].

Further analyses for reconstituted blood T-cell population in nude mice after adult thymus transplant confirmed the presence of circulating CD3+CD4+, CD3+CD8+ T cells as well CD4+FoxP3 Treg cells, but with number of cells matching aged controls. Similarly, CD4/CD8 T-cells present in native and ectopic thymuses also matched aged controls. Finally functional assay with anti-CD3 stimulation, IFNγ expression levels in the media and allogeneic graft skin rejection showed levels of T-cell functions from ectopic thymuses corresponding to the level of aged involuted native thymus.

In summary, we observed that transplanting aging thymuses (up to 8 months old) into a highly vascularized and enriched site as the lymph node was able to successfully rescue immune function in athymic nude mice. However, transplanting very old thymuses (11- and 14-month-old) did not result in any significant rescue in thymic function. Essentially, the ectopic thymuses in these groups behaved similar to the native thymuses of that age.

Our future studies will focus on determining the efficacy of therapeutic interventions such as growth factors and/or cells administration in combination with lymph node transplants to boost thymic epithelial cell proliferation, which may mediate downstream effects of thymus function [36–38]. In addition, it would be interesting to determine whether transplanting a newborn thymus in the lymph node of an aging mouse would rejuvenate immune function in these mice. These studies have tremendous potential to better understand immunologic aging and remedy the age-associated decline in immune function observed in infections and neoplasms.

## Acknowledgments

A.R. designed the study, performed most of the experiments, analyzed the data, prepared the figures, and wrote the manuscript; A.G. performed most of the experiments, analyzed the data, prepared the figures; E.L. designed and supervised the study, and wrote the manuscript; all authors discussed the results and approved the final version of the manuscript.

---

**Figure S1:**
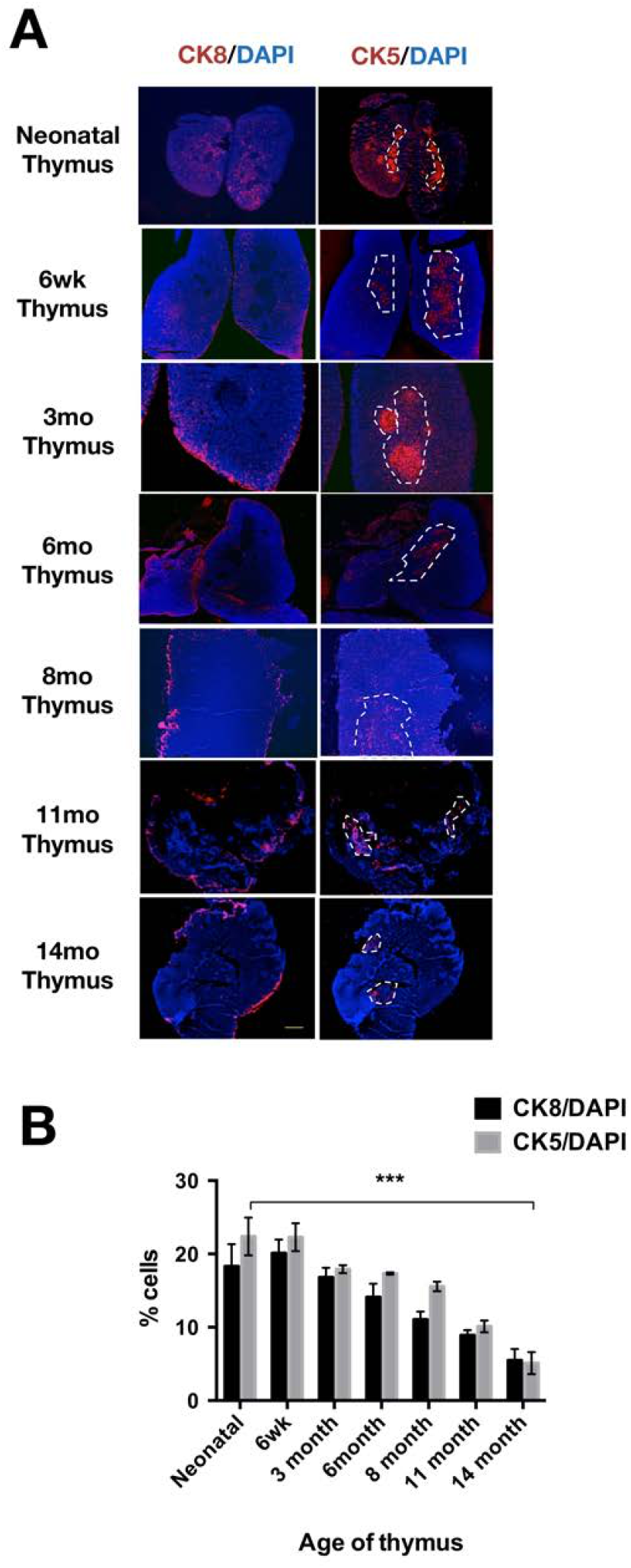
Effect of aging on epithelial cell coverage in thymus. BALB/c thymuses from different aged mice (as indicated) were harvested and frozen in OCT blocks. A) 5-7 μm frozen sections of different aged thymuses were assessed for epithelial cell – cortical epithelial cells (Cytokeratin 8 (CK8), stained red, left panel) and medullary epithelial cells (Cytokeratin 5 (CK5), stained red, right panel). DAPI was used to counterstain nuclei. Scale bar = 50 μm. B) The entire frozen section was imaged using a scanning fluorescence microscope (Nikon 90i) and assessed for various markers. Images were then stitched together and analyzed using the Nikon elements software. Percentage of epithelial cells was calculated using the formula: % cortical/ medullary epithelial cell coverage = (Number of CK5+/CK8+ cells/ Number of DAPI+ cells) x100. **P< 0.005

**Figure S2:**
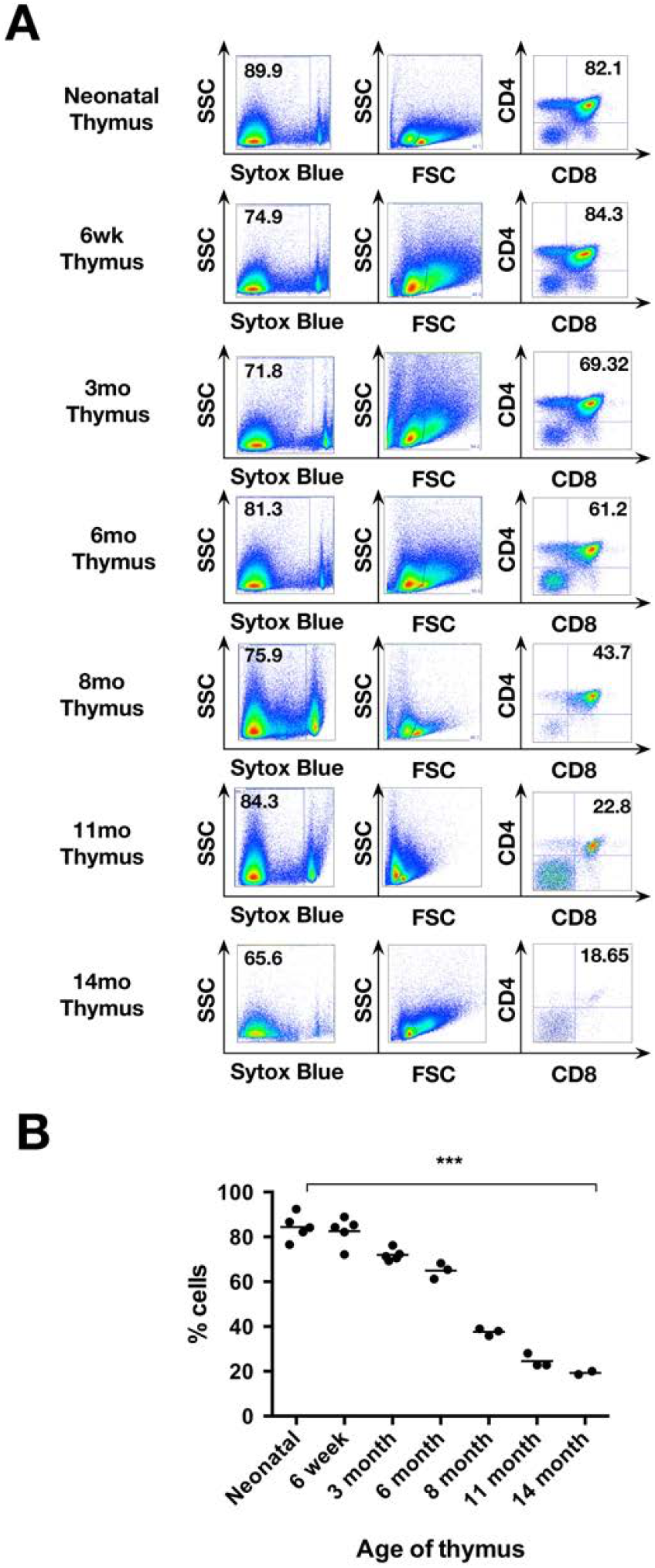
T cell development in mice of different ages. Thymuses from mice of different ages were harvested and dissociated into single cell suspensions. The cells were analyzed for double-positive CD4/CD8 T cells. A) Plots shown are representative of tissues from one mouse from a cohort of 3-5 mice in each group. Gating strategy used was as follows: SSC vs. Live (Sytox Blue) →FSC vs. SSC →CD4 vs. CD8. B) Summary graphs of the proportion of CD4+CD8+ T cells from all the analyzed mice. *** P< 0.0001

**Figure S3:**
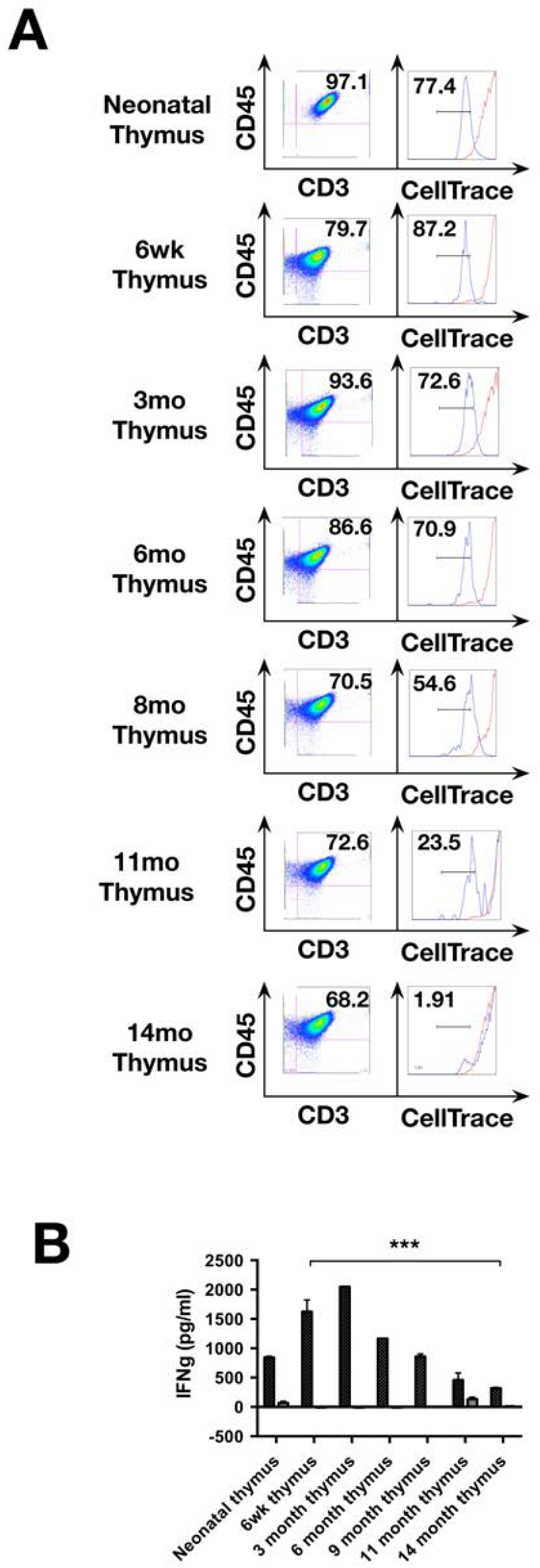
Effect of aging on T cell stimulation. A) Splenocytes were isolated from mice of different ages; CD3+ cells were isolated by MACS and labeled with CellTrace. The cells were then stimulated with anti-CD3 antibody and the cells were assessed for Celltrace dilution (Panel A, blue histograms). Unstimulated cells were used as controls (Panel A, red histograms). B) IFNγ levels from supernatants of cell cultures in (A). ***P<0.0001.

**Table S1:**
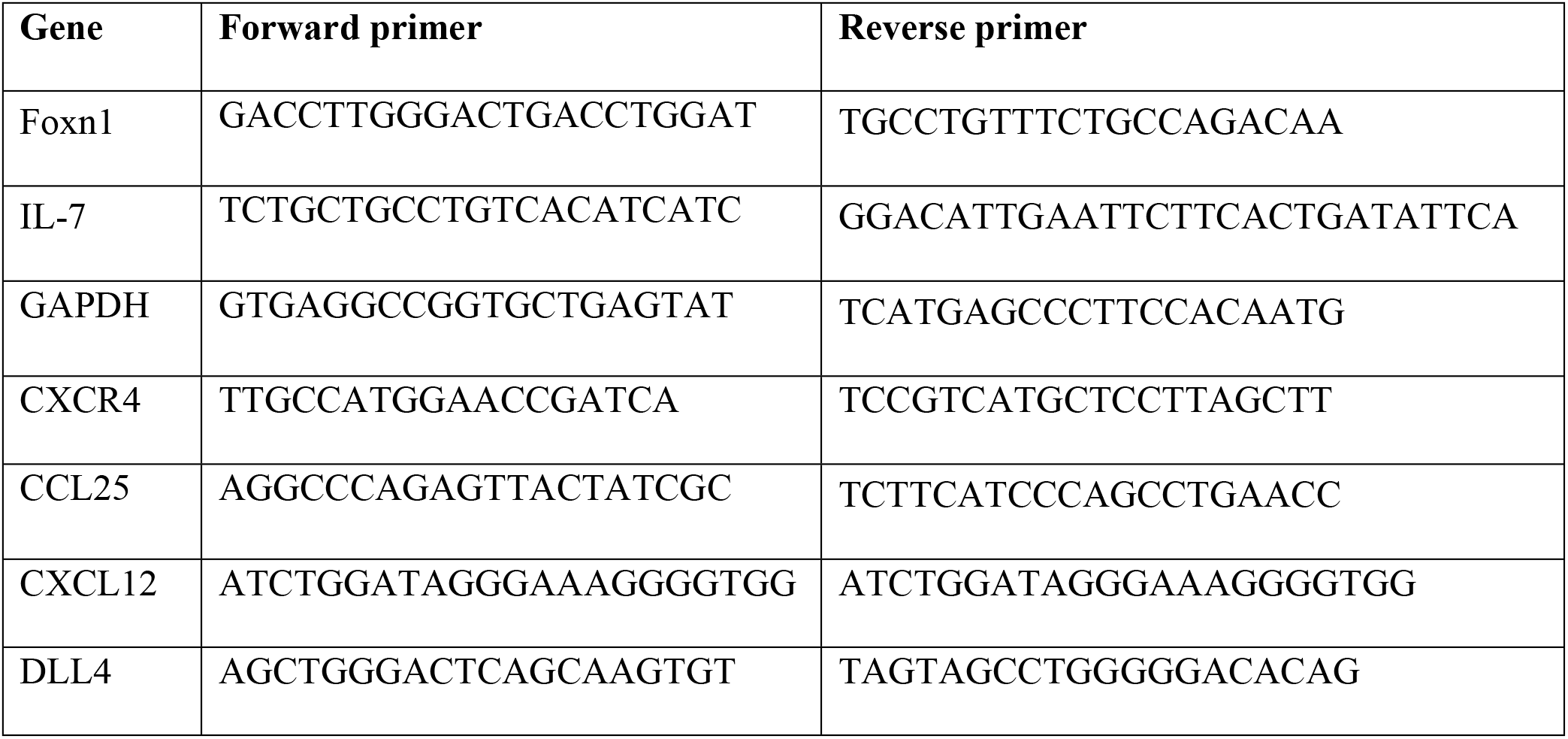
Primer sequences for thymus development genes.

## Notes

### Competing Interest Statement

I have read the journal's policy and one of the authors of this manuscript have the following competing interests: Dr. Lagasse is Chief Scientific Officer of, consults for and owns stock in LyGenesis Inc. Dr. Rao and Ms. Gupta have declared that no competing interests exist.

